# Metabolic Divergence Between Healthy and Conditional Presenilin-1/2 knockout Dementia Model Mice under Chronic Nicotine exposure

**DOI:** 10.1101/2024.11.19.624412

**Authors:** Youwen Si, Shuai Qiu, Lina Wang, Bo Meng, Feiyan Qi

## Abstract

Nicotine, a primary bioactive compound in tobacco, influences a wide range of metabolic pathways, with potential implications for neurodegenerative diseases such as Alzheimer’s disease (AD). This study investigates the metabolic effects of chronic nicotine administration in presenilin-1/2 double knockout (DKO) mice and wild-type (WT) control mice. Eight-month-old DKO and WT mice underwent a three-month oral nicotine administration which did not significantly affect body weight or spontaneous locomotor activity in either DKO or WT mice, as measured by weekly body weight measurement and open field test.

The metabolomic analyses were performed using untargeted LC-MS and GC-MS techniques. The results revealed distinct metabolic profiles between DKO and WT mice under nicotine exposure. In DKO mice, nicotine exacerbated deficits in purine and nicotinamide metabolism, further highlighting dysregulation in energy homeostasis. Key findings include reduced adenosine and elevated dopamine levels in DKO serum, alongside increased cotinine concentrations, suggesting altered nicotine metabolism. WT mice showed significant changes in energy and lipid metabolism pathways, though with less pronounced alterations compared to DKO mice. Enrichment analyses identified significant pathway disruptions, particularly in choline metabolism, amino acid metabolism, and central carbon metabolism, with genotype-dependent responses to nicotine.

This study highlights the intrinsic metabolic alterations in DKO mice, such as disruptions in purine and nicotinamide metabolism, which contribute to their neurodegenerative pathology and unique responses to nicotine. By revealing the differential metabolic impacts of nicotine on healthy and neurodegenerative states, the findings provide valuable insights into its dual therapeutic and pathological roles. These results may guide future research into the metabolic underpinnings of neurodegeneration and the potential for nicotine as a targeted therapeutic intervention.

## Introduction

Nicotine, the primary active compound in tobacco, exerts its effects through the activation of nicotinic cholinergic receptors (nAChRs) distributed throughout both the central nervous system (CNS) and peripheral nervous system (PNS) system ^1–3^. Upon entering the body, nicotine initiates a cascade of physiological responses, significantly impacting various metabolic and cellular signaling pathways, and ultimately, broader physiological and behavioral functions on the organism. In intact animals and humans, nicotine’s physiological effects are well-documented, including an increase in pulse rate, elevated blood pressure, a rise in plasma free fatty acids, mobilization of blood glucose, and heightened catecholamine levels in the bloodstream ^4–7^, has been shown to significantly enhance attention, learning, memory and immune function^8–11^. At the cellular level, nicotine’s interaction with nAChRs triggers a range of downstream signaling pathways that further influence cellular metabolism, neurotransmitter release, and oxidative stress ^7,12–15^. These widespread actions of nicotine underscore its broad impact on the organism’s physiological state.

Given nicotine’s status as a major component of tobacco, its effects have long been a subject of intense debate. On one hand, extensive epidemiological studies have clearly established associations between nicotine exposure (mostly through tobacco smoking) and various diseases such as cancer (especially the lung cancer), cardiovascular disease, metabolic disorders etc., raising significant public health concerns ^16^. On the other hand, nicotine has also been recognized for its potential as a pharmaceutical agent. For instance, there are reports suggesting that nicotine may have beneficial effects in the context of neurodegenerative diseases, such as Alzheimer’s disease (AD) and Parkinson’s disease (PD) etc., highlighting its complex role in human health ^17,18^. The long-standing debate over the effects of nicotine on metabolism and other physiological processes may partly stem from the fact that some studies are based on clinical or epidemiological research involving nicotine intake through tobacco use. Beyond nicotine as the primary component, tobacco contains over 400 other compounds. Therefore, the observed effects on the body following tobacco exposure may be complicated by the presence of other toxic substances, leading to a more complex interaction.

AD, characterized by the progressive deterioration of memory and cognitive function, is the most common neurodegenerative disorder and estimated to affect over 5.8 million individuals in the United States and more than 50 million worldwide, with nearly half of those affected being 75 years or older^19^. There are many hypotheses for AD such as the cholinergic hypothesis, neuroinflammation hypothesis, Aβ deposition hypothesis, Tau protein hypothesis etc., however, the exact pathogenesis is still obscure. The earliest and very important theory of the pathophysiology of AD is the cholinergic hypothesis^20^, which plays a pivotal player in memory processes ^21,22^ and anti-inflammatory functions ^23^. Previous studies have indicated that nicotine can improve cognitive measures in non-smoking patients with mild cognitive impairment (MCI) ^24^, and reduce plaque burden in amyloid precursor protein (APP) transgenic mice ^25–27^. However, these studies did not comprehensively address the impact of nicotine on learning and memory deficiencies or its broader effects on neurodegenerative processes beyond Aβ pathology.

The Aβ hypothesis of AD has faced increasing scrutiny due to the limited success of Aβ-targeted therapies in reducing cerebral Aβ^28^. This has highlighted the importance of exploring Aβ-independent mechanisms in AD pathology. Presenilins (PS), as part of the γ-secretase complex, are critical for the cleavage of APP and Notch receptors and pathogenic mutations in PS genes account for most familial Alzheimer’s disease cases ^29^. Conditional PS1 and PS2 double knockout (DKO) mice obtained by crossing the forebrain-specifically PS1 knocked out mice with systemically PS2 gene knocked out mice, exhibited AD-like symptoms including age-dependent specific forebrain atrophy, tau protein hyperphosphorylation, massive neuronal apoptosis, glial cell proliferation and cognitive dysfunction, but without Aβ deposition as the absence of the functional γ-secretase. Therefore, DKO mice can serve as a unique model for studying Aβ-independent neurodegenerative mechanisms ^30–32^.

In this study, we aimed to investigate the effects of chronic nicotine administration on metabolism in both neurodegenerative disease processes and normal physiological states. We utilized 8-month DKO model mice, which exhibit neurodegenerative pathology, alongside age-matched healthy control mice. Following a three-month regimen of oral chronic nicotine administration, we conducted untargeted metabolomics analysis using LC-MS and GC-MS techniques. By comparing the metabolic profiles of the neurodegenerative model mice to those of the control group, we examined how nicotine treatment differentially impacts these groups. Our findings provide metabolomic evidence for the potential therapeutic application of nicotine in neurodegenerative diseases, highlighting the metabolic alterations associated with nicotine exposure.

## Materials and Methods

### Mice and Nicotine Administration

The DKO mice were generated by crossing forebrain PS1 knockout mice (`a`CaMKII-Cre+, PS1 flox/flox) with conventional PS2 knockout mice (PS2-/-) as previously described ^31^. The littermate WT mice (Cre-, PS1+/+, PS2+/+) served as the control mice. Both DKO and WT mice were maintained on the C57BL6/J background. Eight-month-old DKO and WT mice were randomly divided into nicotine-treated (DKO-N and WT-N, respectively) and blank control (DKO-B and WT-B, respectively) groups with equal proportions of males and females using a computer-generated randomization schedule. The sample number of each group ranges from 9-11.

DKO and WT mice were housed at 24°C and 40-70% humidity with *ad libitum* access to food and water (named as DKO-B and WT-B group, respectively) or water with nicotine (100 μg/ml (-)-nicotine (Alta Scientific, 1ST20302) (named as DKO-N and WT-N group, respectively) and on a 12-hour light/dark cycle (light on from 7:00 am to 7:00 pm) in the ECNU Public Platform for Innovation (010). The nicotine was administrated for 3 months. The water intake was measured by recording the total amount of water consumed by the mice over a one-week period. The average water intake per mouse was then calculated by dividing the total water consumption by the number of mice in each group. The experiments were approved by the Animal Ethics Committee of East China Normal University.

### Open Field Test

The open field test was performed to measure the spontaneous activity of the mice. The mice were handled for 5 min per day for three consecutive days before the behavioral tests to eliminate the nervous emotion of the mice. The open field test was performed using the IR Actimeter software (Panlab). The box size was 22 cm (L) × 22 cm (W) × 38 cm (H). The central zone was defined as a square area of 12 cm in the center. The total movement distance of the locomotor activity was recorded for 5 min and analyzed by the software.

### Samples Collection

Mice were deeply anesthetized with pentobarbital sodium (30 mg/kg). Blood samples were collected from the ocular vasculature and centrifuged at 4000× g for 10 min, and the supernatant was aspirated to obtain serum for metabolomic analysis and cotinine ELISA. Their prefrontal forebrain cortex tissues were quickly removed on ice and then transferred to liquid nitrogen and stored at −80°C for later analysis.

### ELISA Assay

The determination of serum cotinine levels was conducted utilizing the Cotinine ELISA kit (Abnova, KA2264). Briefly, 10 µL of serum samples were aliquoted into respective wells of a 96-well plate, with each sample analyzed in triplicate. Then 100 µl Cotinine HRP Enzyme Conjugate was added, thoroughly mixed, and then incubated for 60 min at room temperature in a light-protected environment. after wash, 100 µl of TMB substrate was added and incubate for 15 minutes. The enzymatic reaction was terminated by the addition of 100 µL of stop solution, followed by thorough mixing. The absorbance of the resulting solution was measured at 450 nm using an enzyme marker (Infinite M Nano, Tecan, Germany). To quantify the cotinine concentration in the samples, a calibration curve was established, and concentrations were calculated using specialized ELISA software.

### Liquid chromatography–mass spectrometry (LC/MS) untargeted metabolome analysis

LC/MS untargeted metabolome analysi**s** was performed by Shanghai Luming biological technology co.Ltd (Shanghai, China). Each group included six samples, except for the DKO-B group, where one sample was excluded during quality control, resulting in a final sample size of five. 100 μL of sample was added to a 1.5 mL Eppendorf tube with 10 μL of 2-chloral-phenylalanine (0.3 mg/mL) dissolved in methanol as internal standard, and the tube was vortexed for 10 sec. Subsequently, 300 μL of an ice-cold mixture of methanol and acetonitrile (2/1, v/v) was added, and the mixtures were vortexed for 1 min, ultrasonicated at ambient temperature (25□ to 28□) for 10 min, stored at −20□ for 30 min. The extract was centrifuged at 13000 rpm, 4□ for 10 min. 300 μL of supernatant in a brown and glass vial was dried in a freeze concentration centrifugal dryer. 400 μL mixtures of methanol and water (1/4, vol/vol) were added to each sample, samples vortexed for 30 sec and ultrasonicated for 3 min, then placed at −20□ for 2 hours. Samples were centrifuged at 13000 rpm, 4□ for 10 min. The supernatants (150 μL) from each tube were collected using crystal syringes, filtered through 0.22 μm microfilters and transferred to LC vials. The vials were stored at −80°C until LC-MS analysis. QC samples were prepared by mixing aliquots of all samples to be a pooled sample.

An ACQUITY UHPLC system (Waters Corporation, Milford, USA) coupled with an QE plus (Thermo Scientific) was used to analyze the metabolic profiling in both ESI positive and ESI negative ion modes. An ACQUITY UPLC HSS T3 column (100 mm × 2.1 mm, 1.8 μm) were employed in both positive and negative modes. The acquired LC-MS raw data were analyzed by the progqenesis QI software (Waters Corporation, Milford, USA). The differential metabolites were selected on the basis of the combination of a statistically significant threshold of variable influence on projection (VIP) values obtained from the OPLS-DA model and *p* values from a two-tailed Student’s *t* test on the normalized peak areas, where metabolites with VIP values larger than 1.0 and p values less than 0.05 were considered as differential metabolites.

### Gas chromatography-mass spectrometry (GC-MS) untargeted metabolome analysis

GS/MS untargeted metabolome analysis was performed by Shanghai Luming biological technology co.Ltd (Shanghai, China). Each group included six samples, except for the DKO-B group, where one sample was excluded during quality control, resulting in a final sample size of five. 50 μL of sample was added to a 1.5 mL Eppendorf tube with 10 μL of 2-chloral-phenylalanine (0.3 mg/mL) dissolved in methanol as internal standard, and the tube was vortexed for 10 sec. Subsequently, an ice-cold mixture of methanol and acetonitrile (2/1, v/v) was added, and the mixtures were vortexed for 30 sec, ultrasonicated at ice-cold temperature for 10 min, stored at −20□ for 30 min. The extract was centrifuged at 13000 rpm, 4□ for 10 min. An aliquot of the 150 μL supernatant was transferred to a glass sampling vial for vacuum-dry at room temperature. And 80 μL of 15 mg/mL methoxylamine hydrochloride in pyridine was subsequently added. The resultant mixture was vortexed vigorously for 2 min and incubated at 37□ for 90 min. 80 μL of BSTFA (with 1% TMCS) and 20 μL n-hexane was added into the mixture, which was vortexed vigorously for 2 min and then derivatized at 70□ for 60 min. The samples were placed at ambient temperature for 30 min before GC-MS analysis. The GC-MS analysis process includes sample pretreatment, metabolite extraction, metabolite derivatization, GCMS detection, data pre-processing and statistical analysis. The derivatived samples were analyzed on an Agilent 7890B gas chromatography system coupled to an Agilent 5977A MSD system (Agilent Technologies Inc., CA, USA). A DB-5MS fused-silica capillary column (30 m × 0.25 mm × 0.25 μm, Agilent J & W Scientific, Folsom, CA, USA) was utilized to separate the derivatives. AnalysisBaseFileConverter software was used to convert the raw data (.D format) to .abf format, and then the .abf data were imported into the MD-DIAL software for data processing. The differential metabolites were selected on the basis of the combination of a statistically significant threshold of variable influence on projection (VIP) values obtained from the OPLS-DA model and p values from a two-tailed Student’s t-test on the normalized peak areas from different groups, where metabolites with VIP values larger than 1.0 and p values less than 0.05 were considered as differential metabolites.

### Statistical analysis

The data is expressed as the mean ± standard error of the mean (SEM). Statistical analysis was performed using repeated measure ANOVA, Student’s *t*-test or one-way analysis of variance (ANOVA) followed by Tukey’s multiple comparison test according to the experimental design with GraphPad Prism software. Significance levels are indicated as follows: \**P* < 0.05, \*\**P* < 0.01, and \*\*\**P* < 0.001.

## RESULTS

### DKO mice exhibited sustained hyperactivity following oral nicotine administration

DKO mice were obtained by crossing the forebrain-specifically (`a`CaMKII-Cre+, PS1 flox/flox) PS1 knocked out mice with systemically PS2 gene knocked out mice. According to our previous study, DKO mice showed age-dependent progressive neurodegeneration, and DKO mice older than 6 months showed a serial of behavior abnormal including sever learning and memory deficits as well as forebrain atrophy, brain inflammation, glial cells activation, tau protein hyperphosphorylation, etc., mimicking the late phase of AD^32,33^. In this study, 8-month-old DKO and WT mice were used, and after chronic nicotine administration by drinking water for 3 months at a dosage of 100 μg/mL.

Epidemiological studies have generally found that long-term smoking is associated with lower body weight compared to non-smokers. Nicotine can increase metabolism, leading to reduced caloric intake and higher energy expenditure^34^. In our experiments, the drinking consumption was measured for 12 weeks (the measurement stopped when the behavior tests began) to exclude the possible impact on water intake and consumption by adding nicotine. The results showed no significant difference among the four groups during the nicotine administration (data not shown). Before the nicotine exposure, the body weight of the DKO and WT group of mice showed no difference. During the nicotine administration, the body weight was measured once a week and the overall statistical analysis reveals a significant effect of time (Week) on body weight of both male and female mice (using Greenhouse-Geisser correction, for male mice, *F* _(1.492, 22.376)_ = 187.063, P < 0.001; for female mice, *F* _(2.995, 53.915)_ = 143.956, P < 0.001;) but no significant effect of group, (for male, *F* _(3, 15)_ = 0.468, P = 0.709; for female, *F* _(3, 18)_ = 0.155, P = 0.925). Additionally, no significant interaction between time and group was found (using Greenhouse-Geisser correction*, F* _(4.475, 22.376)_ = 1.000, P = 0.434 for male; and *F* _(8.986, 53.915)_ = 0.998, P = 0.453 for female, respectively), suggesting that while body weight changed significantly over time, the patterns of weight change did not differ between the groups (Figure 1A). Therefore, nicotine in the dose of 100 μg/ml has no significant effect on the body weight of both WT and DKO mice. All these results eliminated the possibility of differences in nicotine intake among the four groups of mice, and nicotine has no obvious effect on the body weight increase with time.

**Figure 1.**
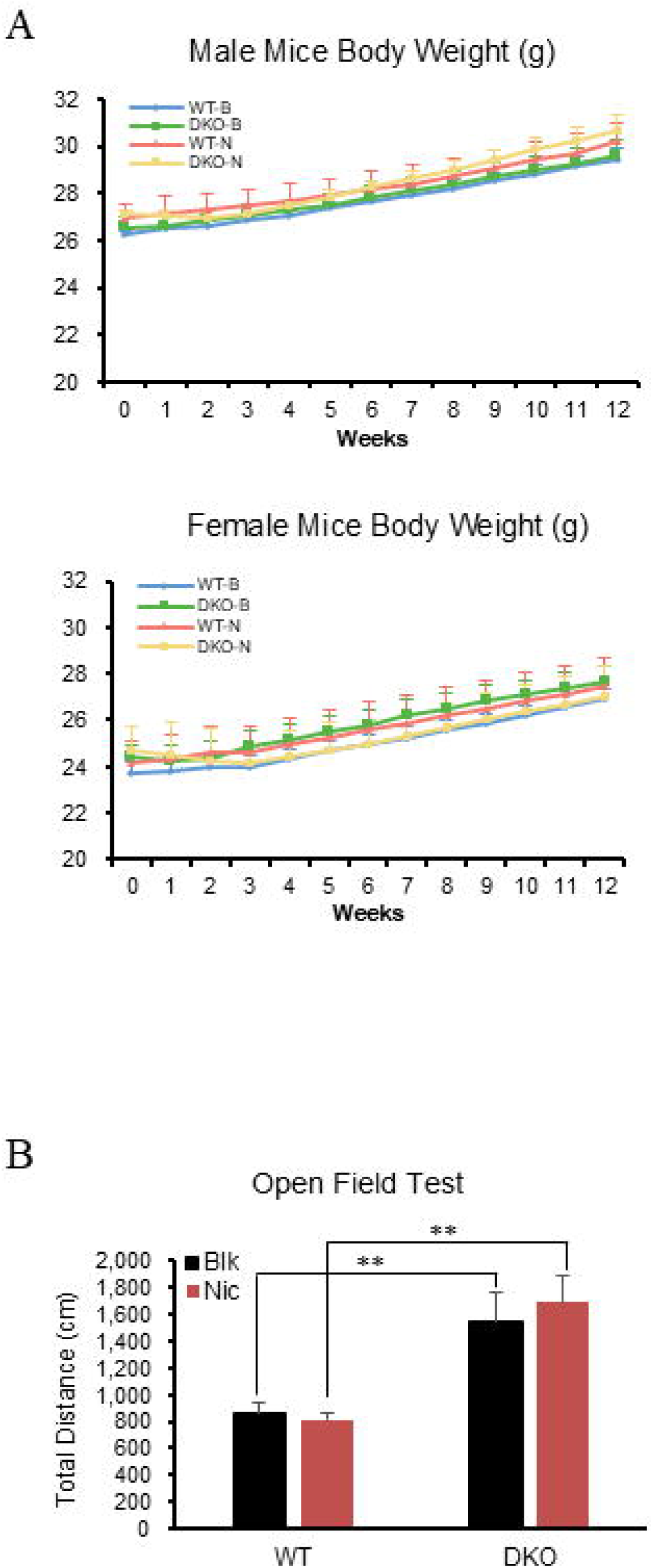
(A) Body weight measurement. The body weight of male (left) and female (right) mice of the four groups were measured once a week during the experiment. (B) The open field test. The total movement distance (left) and locomotion activities were measured in 5 min. WT-B: WT mice with normal drinking water (n=9); WT-N: WT mice treated with nicotine in drinking water (n=11). DKO-B: DKO mice with normal drinking water (n=9); DKO-N: DKO mice with nicotine in drinking water (n=9). The results are presented as mean ± S.E.M. \**P* <0.05, \*\**P* < 0.01, and N.S. represents no significant difference.

After 3-month nicotine administration, the open field test was performed for the spontaneous activity measurement on the mice at approximately 11 months of age. As shown in Figure 1B, the total movement distance and locomotion activities of DKO mice were significantly higher than those of WT mice, resulting almost double distance difference, which is consistent with our previous results^32^, and the difference of spontaneous activity among the mice didn’t result in significant variation in body weight. Multivariate analysis results showed that genotype had a significant effect on movement distance in the open field test, as indicated by Pillai’s Trace, *F*_(2, 74)_ = 15.134, p < 0.001; nicotine treatment had no significant effect (*F*_(2, 74)_ = 0.089, p = 0.915, and the interaction between genotype and treatment (genotype × treatment) was not significant ( *F*_(2, 74)_ = 0.283, p = 0.755). These results indicate that the effect of nicotine treatment on the spontaneous activity in both WT mice and DKO mice was consistent across genotypes, with no differential responses based on genotype.

### Distinct Effect of Nicotine on the Metabolism Profile of DKO and WT Mice by LC□MS

Following oral administration, nicotine is absorbed into the body and undergoes a series of metabolic reactions, resulting in the production of metabolites such as cotinine^35^. Cotinine is commonly used as a biomarker to assess the rate of nicotine absorption in the body^36^. The Elisa result showed that DKO mice had higher serum cotinine concentrations than WT mice after nicotine administration via drinking water by ELISA (WT-N: 76.98 ± 38.02 ng/ml, DKO-N: 636 ± 270.5 ng/ml, *P* value = 0.05) (Figure 2A). In the serum of DKO-B or WT-B mice, the cotinine content was below the detectable range (data not shown).

**Figure 2.**
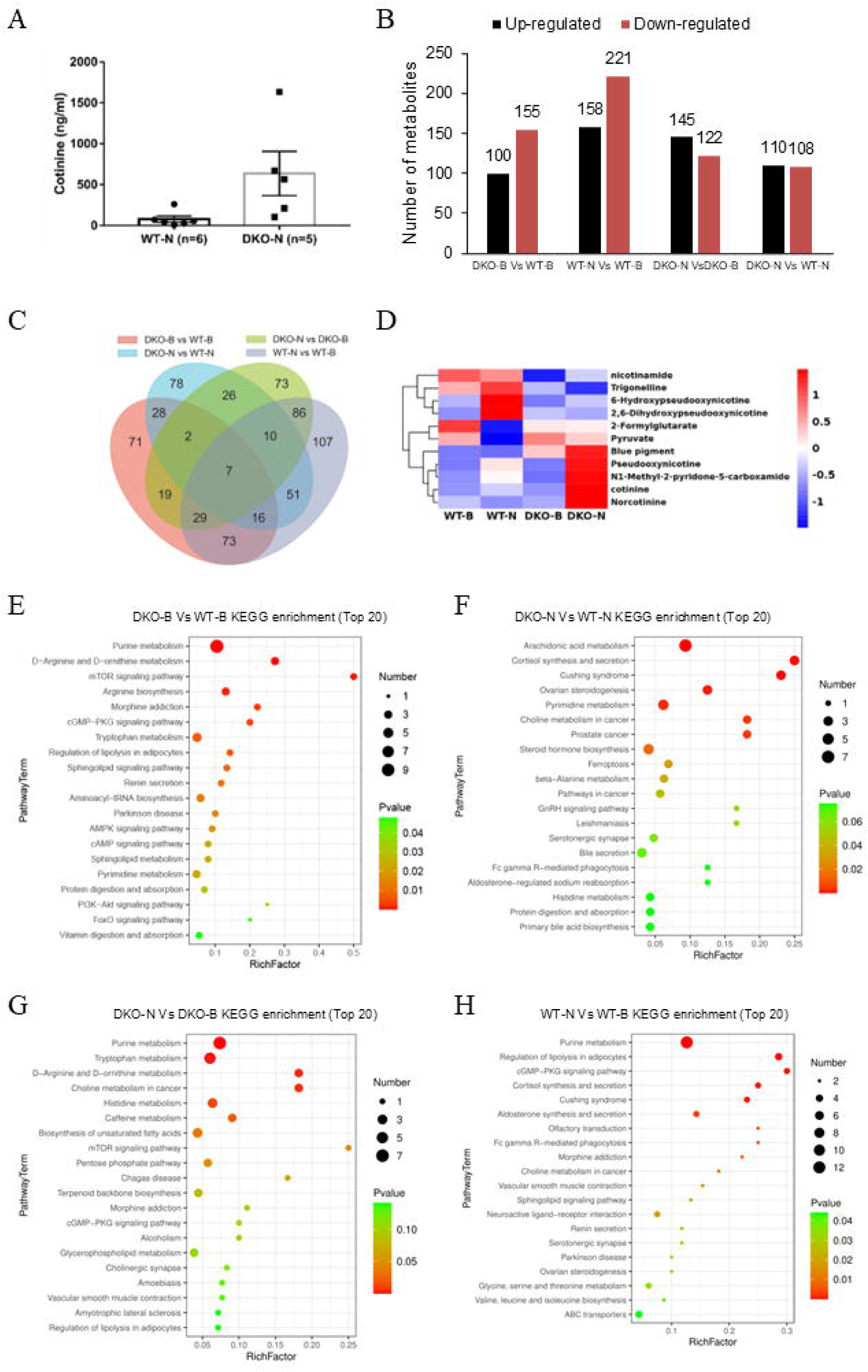
(A) Cotinine level in the serum of the DKO-N and WT-N mice measured by ELISA. (B-H) Untargeted LC□MS metabolomics analysis. (B) up-regulated and down-regulated SCMs in the four comparative groups. (C) Venn diagram showing the numbers of SCMs unique to each of the four comparative groups and those shared among them. (D) Hierarchical clustering of SCMs in the intersection of four comparative groups. (E-H) Bubble diagram of the top 20 differential metabolic pathways in the four comparative groups.

Metabolites are downstream of transcriptional and proteomic changes; they can provide instantaneous “snapshots” of the state of a cell or organism. Both LC-MS and GC-MS are wildly employed to analyze metabolome, and these two techniques complement each other in terms of the types of metabolites. LC-MS is particularly suited for identifying polar and non-volatile metabolites, such as amino acids, peptides, sugars, lipids, and nucleotides. In contrast, GC-MS excels in detecting volatile and low molecular weight metabolites, including fatty acids, alcohols, ketones, and small organic acids, especially after derivatization. By utilizing both techniques, we aimed to achieve a comprehensive coverage of the metabolome, capturing both volatile and non-volatile compounds in the serum.

Firstly, the relative concentrations of differentially abundant metabolites in WT and DKO mice under nicotine conditions were measured by LC-MS (Supplementary table 1). Principal component analysis (PCA) plot results showed a trend of separation among the four groups of samples with good analytical stability and experimental reproducibility (Supplementary Figure S1).

We established four comparison groups: DKO-B vs WT-B, WT-N vs WT-B, DKO-N vs DKO-B, and DKO-N vs WT-N. With nicotine exposure, there were more downregulated significantly changed metabolites (SCMs) than upregulated SCMs in WT mice and DKO mice (Figure 2B). The number of SCMs that were unique or shared between these four comparison groups is shown in the Venn diagram (Figure 2C). And figure 2E showed the hierarchical clustering of SCMs in the intersection of four comparative groups. Particularly, Adenosine content was significantly decreased in DKO mice compared to WT mice, reaching only 23±11% of the adenosine level observed in WT mice (supplemental figure S2) and nicotine exposure led to a near-complete depletion of adenosine in the serum of both DKO and WT mice, reducing levels to approximately 1% of those observed without nicotine exposure. The dopamine levels in the serum of DKO mice were significantly elevated compared to the WT group, and nicotine exposure did not significantly alter serum dopamine levels in either DKO or WT mice. Notably, nicotine metabolites, including cotinine and norcotinine, were increased higher in DKO mice than those in WT mice after nicotine administration, which are consistent with the cotinine’ ELISA result and further confirmed a dysregulated nicotine metabolism in DKO mice.

We generalized all SCMs for pathway enrichment according to the KEGG database. The top 20 affected metabolic pathways are shown in Figure 2E-H by *P* value scoring. Firstly, we compared the differential metabolites between DKO and WT mice without nicotine exposure and found that the primary SCMs were related to purine metabolism, amino acid metabolism, mTOR signaling pathway, morphine addiction, lipolysis, etc. Most of the KEGG pathways enriched in the “DKO-B vs WT-B” group were altered after nicotine administration. The top enriched KEGG pathways of “DKO-N vs WT-N” were arachidonic acid metabolism, hormones synthesis and secretion, choline metabolism and serotonergic synapse, etc. There were 38 shared and 54 unique metabolic pathways between the WT-N vs WT-B and DKO-N vs DKO-B groups. Purine metabolism was significantly impacted in the DKO group, similar to the WT mice; “Choline metabolism” and “cholinergic synapse” were also enriched in both the “WT-N vs WT-B” and “DKO-N vs DKO-B” groups, while they showed marked changes in the DKO mice, whereas were less pronounced in the WT group with nicotine exposure, which further confirmed that nicotine affects the choline metabolism in both WT and DKO mice but to different extents..

### Distinct Effect of Nicotine on the Metabolism Profile in DKO and WT Mice by GC-MS

The metabolism profile of the serum in DKO and WT mice with/without nicotine administration was measured by GC-MS untargeted metabolome analysis (Supplementary table 2), and the results revealed a total of 34 significantly dysregulated metabolites in DKO-B mice compared to WT-B mice; after nicotine administration, there were 41 dysregulated metabolites in DKO mice and 37 in WT mice (Figure 3A), and figure 3B showed the hierarchical clustering of SCMs in the four comparative groups. Similar to LC/MS, the metabolic difference between DKO and WT changed significantly with nicotine administration, with only 4 metabolites remaining with/without nicotine exposure. Among them, the nicotinamide content was significantly reduced in the serum of DKO mice, reaching 60 ± 16% of the levels observed in WT mice, and nicotine exposure didn’t significantly affect its level in either genotype (supplemental figure S3).

**Figure 3.**
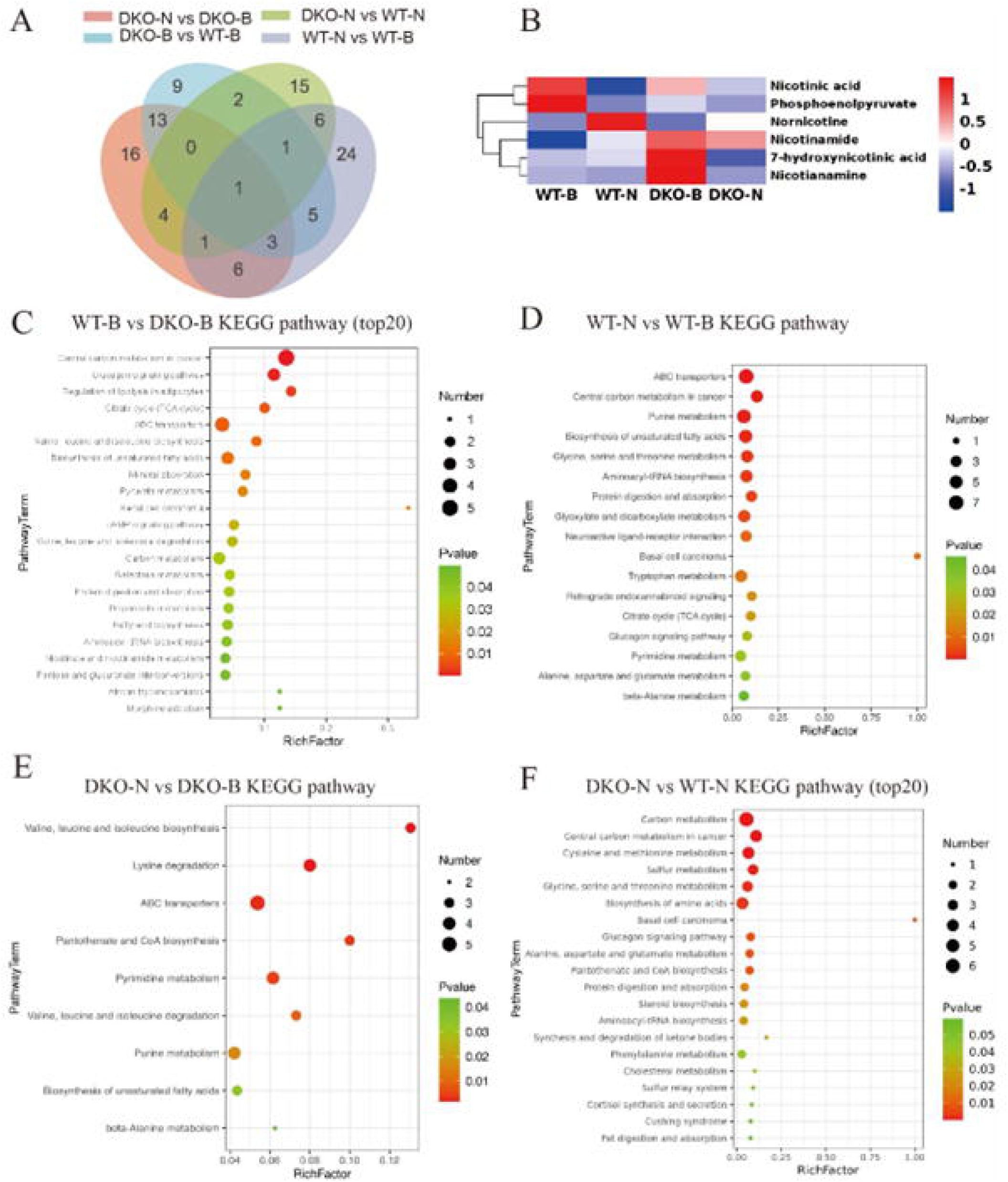
Untargeted GC/MS metabolome analysis. (A) Venn diagram showing the numbers of SCMs unique to each of the four comparative groups and those shared among them. (B) Hierarchical clustering of SCMs in the four comparative groups. (C-F) KEGG pathways of differentially abundant metabolites of the groups.

KEGG functional enrichment analyses were also performed for functional prediction of differentially abundant metabolites. Based on the negative log(P) and impact values, the most enriched KEGG term was central carbon metabolism when we compared DKO mice with WT mice, followed by glucagon signaling, regulation of lipolysis in adipocytes and the citrate cycle, which all indicated dysregulated energy metabolism in DKO mice. Similar to the LC/MS results, the nicotinate and nicotinamide metabolism signaling pathway was also enriched, indicating impaired nicotinate and nicotinamide metabolism in DKO mice (Figure 3C).

Different KEGG pathways of differentially abundant metabolites were enriched in WT mice and DKO mice exposed to nicotine. With nicotine exposure in WT mice, the most enriched KEGG pathway of SCMs was energy metabolism, purine metabolism, and the synthesis of unsaturated fatty acids, as well as protein digestion and absorption. (Figure 3D); however, nicotine administration in DKO mice predominantly affected various amino acid metabolic pathways, and the impact on carbon metabolism was relatively smaller compared to WT mice. (Figure 3E). The primary metabolic differences between DKO and WT mice after nicotine treatment were carbon metabolism pathways, though other pathways also showed changes warranting further investigation.

Both the LC-MS and GC-MS results revealed a systemic metabolic change when PS1 and PS2 genes loss of function in forebrain excitatory neurons in mice, and the chronic nicotine differentially affect the metabolism profiles in DKO and WT mice.

## Discussion

In the pathological progression of neurodegenerative diseases, such as AD, there has been extensive research and numerous reports on changes at the transcriptomic and proteomic levels. In our previous research on DKO dementia model mice, we conducted multi-level approaches using cDNA microarrays, protein microarrays, and RNA sequencing and these revealed the roles of key signaling pathways and inflammatory factors and non-coding RNAs profile changes in the progression of neurodegeneration^37–39^. However, studies focusing on metabolomics are relatively scarce. Investigating the metabolic changes in neurodegenerative diseases is crucial because metabolism is intricately linked to cellular function and energy homeostasis, which are often disrupted in these conditions. We observed significant metabolic differences between the peripheral serum samples of DKO and WT mice, suggesting that neurodegenerative changes not only affect the CNS but also influence the overall metabolic state of the organism through neuro-endocrine interactions. We found a marked decrease in nicotinamide and adenosine content in the DKO mice. Nicotinamide has been proven to provide neuroprotective effects against various toxic conditions, including oxidative stress, hypoxia, excitotoxicity, ethanol-induced neuronal injury, Aß toxicity, age-related vascular diseases, mitochondrial dysfunction, insulin resistance, excessive lactate production, and pancreatic glucose homeostasis loss due to β-cell dysfunction ^40–42^. Therefore, the reduction in nicotinamide levels in DKO mice may be one of the mechanisms underlying their neurodegenerative pathology. It’s notable that DKO mice with nicotine exposure showed a trend of over 20% elevation of nicotinamide level, no statistical significance though, may suggest a beneficial effect. Adenosine, a neuromodulator that is widely distributed across various organs, including the CNS, exerts diverse physiological effects primarily through adenosine receptors. Peripherally, adenosine acts as an immune-suppressant, while in the CNS, it modulates the activity of excitatory neurons through its receptors, thereby playing a role in the initiation of sleep and regulation of the sleep-wake cycle ^43–45^. A decrease in adenosine levels can lead to increased wakefulness and alertness, and potentially resulting in symptoms of insomnia. Previously, we showed the increased wakefulness periods and decreased total time spent in both rapid eye movement (REM) and nonrapid eye movement (NREM) sleep in 6-month-old DKO mice^46^. Here the GC-MS results provide a possible explanation for the circadian rhythm sleep disorder in DKO mice. The direct reduction in adenosine levels in DKO mice also impacts their neurological functions. The nicotine exposure failed to increase the adenosine content in DKO mice, and also dramatically decreased its level in WT mice, which suggests that chronic nicotine exposure may affect the sleep in healthy people. Previously, we measured monoamine neurotransmitter levels in the forebrain cortex and hippocampus of different ages of DKO mice, and found the dynamic profile changes, among which, dopamine level in DKO mice significantly decreased in the forebrain cortex and slightly decreased in the hippocampus at the age of 6 months, while significantly increased in the hippocampus, but no difference in cortex at the age of 12 month^47^. Here we found that the dopamine levels in the serum of DKO mice were significantly elevated compared to the WT group at the age of about 11 months, which further suggests that dopamine levels vary dynamically across different brain regions and between central and peripheral systems during the progression of neurodegeneration.

Nicotine’s effects on the metabolomes of DKO and WT mice revealed substantial differences, particularly within carbon metabolism pathways. These differences may be attributed to several potential mechanisms: in response to nicotine exposure, DKO mice with neurodegenerative alterations display a unique metabolic response, resulting in distinct regulation of systemic metabolic pathways. Additionally, nicotine’s impact on the metabolism of peripheral organs could influence central nervous system activity through the gut-brain axis or the neuro-immune system, further contributing to modifications in central-peripheral regulatory dynamics.

Many people are exposed to nicotine over long periods through smoking or other tobacco products. Long-term trials better simulate the chronic exposure experienced by these individuals, providing more applicable data for public health recommendations and regulations. However, researchers have observed various inconsistent outcomes following nicotine exposure across different animal models. The response to nicotine can also vary significantly based on dose, frequency, mode of administration and stages of AD highlight the complexity of its action on human physiology (Table 1). For example, Sultana et al reported that nicotine improves memory at a low dose and has no effect at a higher dose in ApoE-knockout mice ^48^. Yang et al reported that very low-dose nicotine (2 ug/ml) rebalanced NAD^+^ homeostasis and ameliorated cellular energy metabolism disorders without activation of brain nAChR in aged mice^49^, which further underscore the critical importance of determining the appropriate timing and dosage of nicotine administration when considering its potential as a therapeutic intervention.

In summary, our metabolomic studies on DKO mice suggest that due to alterations in their intrinsic metabolic state, patients with neurodegenerative diseases may exhibit significant differences in drug metabolism compared to healthy individuals. The results also provide insights into the biochemical alterations and metabolic pathways that are affected in neuro-degenerative pathology, potentially revealing new biomarkers for early diagnosis, disease progression, and therapeutic targets. Our research has provided significant insights into nicotine’s mechanisms of action; the nicotine treatment induces significant metabolic changes, impacting various biological pathways, including energy metabolism, lipid metabolism, amino acid metabolism, and neurotransmitter synthesis, and several gaps remain, particularly regarding the differential effects in cognitively normal and cognitively impaired individuals. Future research should address these gaps to further our understanding and optimize nicotine’s therapeutic potential.

## Ethics in publishing

All experiments were approved by the Institutional Animal Care and Use Committee of the East China Normal University (IACUC approval ID #M10020).

## Acknowledgement

This work was financially supported by the grant from the Joint Institute of Tobacco and Health (No.2022539200340036).

## Datasets/Data Availability Statement

The LC□MS datasets examined in this study are accessible in MetaboLights as MTBLS5625 (https://www.ebi.ac.uk/metabolights/MTBLS5625). The GC□MS datasets are available in MetaboLights as MTBLS5627 (https://www.ebi.ac.uk/metabolights/MTBLS5627).

## Author Contributions

Youwen Si (Methodology; Data curation; Writing - Original draft preparation); Shuai Qiu (Data curation); Lina Wang (Resources; Project administration); Bo Meng (Methodology; Writing - Review & Editing; Supervision; Project administration); Feiyan Qi (Conceptualization; Project administration);

## Declaration of conflicting interests

It is important to note that the financial sponsor had no role in the study design, data collection and analysis, publication decision, or manuscript preparation. The decision to publish and the content of the manuscript are solely the responsibility of the authors.

Feiyan Qi is an employee of the Joint Institute of Tobacco and Health. Her contribution to this study was limited to providing general advice on research direction. She was not involved in the design, execution, data analysis, or interpretation of the experiments, nor did she influence the preparation of the manuscript. The authors declare that no conflicts of interest influenced the outcomes or conclusions of this study.

All other authors have no conflict of interest to report.

## Notes

https://www.ebi.ac.uk/metabolights/MTBLS5625

https://www.ebi.ac.uk/metabolights/MTBLS5627

