## Supplementary material for "Metabolic Divergence Between Healthy and Conditional Presenilin-1/2 knockout Dementia Model Mice under Chronic Nicotine exposure": Figures: Supplementary figures and legends.docx

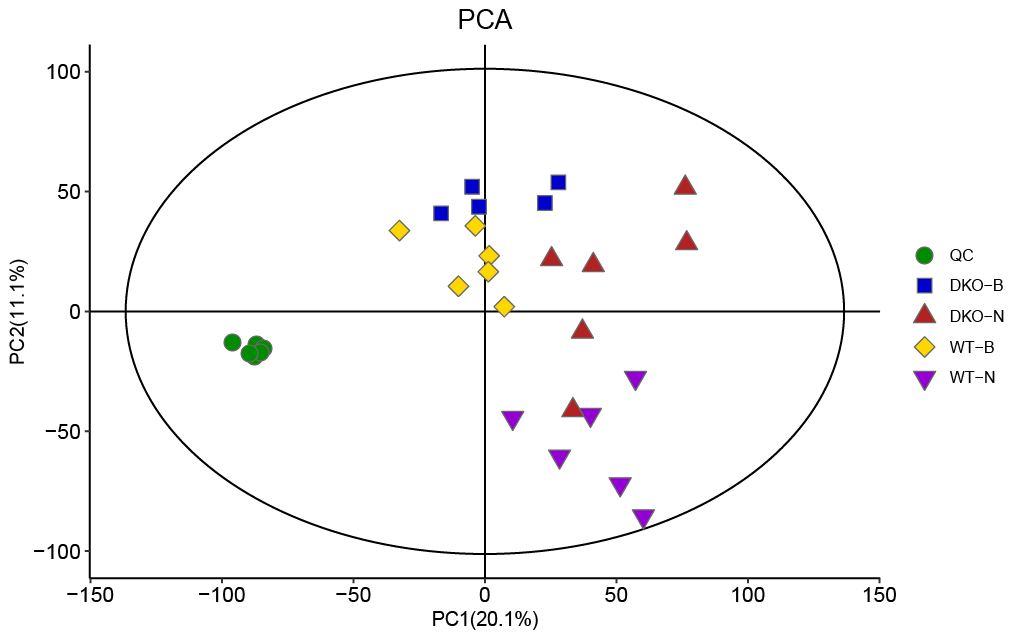

Figure S1. Principal component analysis (PCA) score plots of significantly changed metabolites (SCMs) in both Quality Control (QC) and four treatments in LC-MS assay.

Adenosine

|  | Percentage (%) |
| --- | --- |
| WT-B | 100% |
| DKO-B | 23.1052% |

Figure S2. Adenosine levels in the DKO and WT mice with and without nicotine administration measured by LC-MS.

Nicotinamide

Percentage (%）

|  | WT | DKO |
| --- | --- | --- |
| Blk | 100% | 60.0509% |
| Nic | 93.7967% | 75.8914% |

Figure S3. Nicotinamide levels in the DKO and WT mice with and without nicotine administration measured by GC-MS.
